## Supplementary Materials for "Resilience of small intestinal beneficial bacteria to the toxicity of soybean oil fatty acids"

**Figure 4 – Source data 1.** **SBO mouse diets.**

| Ingredient | 16% SBO diet  (TD.130215) (g/kg) | 44% SBO diet  (TD.130214) (g/kg) |
| --- | --- | --- |
| Casein (ethanol washed),  "Vitamin-Free" Test | 200.0 | 240.0 |
| L-Cystine | 3.0 | 3.6 |
| Corn Starch | 371.986 | 162.354 |
| Maltodextrin | 80.0 | 80.0 |
| Sucrose | 200.0 | 200.0 |
| Soybean Oil | 70.0 | 230.0 |
| Cellulose | 30.0 | 30.0 |
| Mineral Mix, AIN-93G-MX (94046) | 35.0 | 42.0 |
| Vitamin Mix, Teklad (40060) | 10.0 | 12.0 |
| TBHQ, antioxidant | 0.014 | 0.046 |
