## Supplementary Materials for "Resilience of small intestinal beneficial bacteria to the toxicity of soybean oil fatty acids"

**Figure 3 – Source data 1.** **Oligos used to generate *L. reuteri* mutants**

| Recombineering oligos* | | | Screening method |
| --- | --- | --- | --- |
| SDR21 | FabT LR2 deletion | tctgctacttgagaaattgtttgatggttatacatcgtaattgcattaattgttaaatcattgaatgtacttttccgtaactcattttct | Restriction digest with MfeI |
| SDR42 | FabT LR5 SNP with surrounding synonymous mutations | accatcatgttatgaaaggcccgatgtgcacgatatagtacTcTCTcCcGtttggttaaacgtaaccgaacaatgcggcgatcatcttta | MAMA-PCR with SDR44 |
| SDR49 | hydrolase LR5 SNP with surrounding synonymous mutations | agcccagccagctgttcaaaagctagcgaactatactcaTcaCcaCcaTAGCacccgcctgcaacattctattgcggtttcgtatgatag | MAMA-PCR with SDR51 |
| oJP577 | RpoB mutation | tcaaaccaccaggaccaagcgctgaaagacgacgcttTCTGCttaattcacctaatgggttggtttgatccatgaactgg | 25 µg/ml rifampicin |
| Sequencing and MAMA-PCR oligos | | | Annealing temperature |
| SDR13 | FabT forward primer | TCGGGGCTATAGAATAAAATTGAAGGG | 56 |
| SDR14 | FabT reverse primer | CAATGTTTGTCAATCCTTATCTCTAGG | 56 |
| SDR15 | hydrolase forward primer | CTCGTTAAAATCGTGTATGATAATTGC | 58.5 |
| SDR16 | hydrolase reverse primer | CCATAATTTAACTCTCCTTCTCCTC | 58.5 |
| SDR44 | FabT LR5 MAMA-PCR reverse primer | ACGATATAGTACTCTCTCCCG | 56 |
| SDR51 | hydrolase LR5 MAMA-PCR forward primer | GAACTATACTCATCACCACCATAGC | 58.5 |

*Mutations generated by oligo are in caps; for SDR21, the bases adjacent to the created deletion are underlined.
